# Geometric effects in gas vesicle buckling under ultrasound

**DOI:** 10.1101/2022.06.27.497663

**Authors:** Hossein Salahshoor, Yuxing Yao, Przemysław Dutka, Nivin N. Nyström, Zhiyang Jin, Ellen Min, Dina Malounda, Grant J. Jensen, Michael Ortiz, Mikhail G. Shapiro

## Abstract

Acoustic reporter genes based on gas vesicles (GVs) have enabled the use of ultrasound to noninvasively visualize cellular function *in vivo*. The specific detection of GV signals relative to background acoustic scattering in tissues is facilitated by nonlinear ultrasound imaging techniques taking advantage of the sonomechanical buckling of GVs. However, the effect of geometry on the buckling behavior of GVs under exposure to ultrasound has not been studied. To understand such geometric effects, we developed computational models of GVs of various lengths and diameters and used finite element simulations to predict their threshold buckling pressures and post-buckling deformations. We demonstrated that the GV diameter has an inverse cubic relation to the threshold buckling pressure, whereas length has no substantial effect. To complement these simulations, we experimentally probed the effect of geometry on the mechanical properties of GVs and the corresponding nonlinear ultrasound signals. The results of these experiments corroborate our computational predictions. This study provides fundamental insights into how geometry affects the sonomechanical properties of GVs, which, in turn, can inform further engineering of these nanostructures for high-contrast, nonlinear ultrasound imaging.

**STATEMENT OF SIGNIFICANCE:** Gas vesicles (GVs) are an emerging class of genetically encodable and engineerable imaging agents for ultrasound whose sonomechanical buckling generates nonlinear contrast to enable sensitive and specific imaging in highly scattering biological systems. Though the effect of protein composition on GV buckling has been studied, the effect of geometry has not previously been addressed. This study reveals that geometry, especially GV diameter, significantly alters the threshold acoustic pressures required to induce GV buckling. Our computational predictions and experimental results provide fundamental understanding of the relationship between GV geometry and buckling properties and underscore the utility of GVs for nonlinear ultrasound imaging. Additionally, our results provide suggestions to further engineer GVs to enable *in vivo* ultrasound imaging with greater sensitivity and higher contrast.

## INTRODUCTION

Ultrasound imaging has demonstrated tremendous potential for monitoring biological processes due to its deep tissue penetration and non-invasive operation. Recently, the gas vesicle (GV) – a unique genetically encoded, gas-filled, protein-shelled nanostructure – was developed as a new type of contrast agent (1, 2), reporter gene (3–5) and biosensor (6) to connect ultrasound images to dynamic biological activities such as gene expression and enzyme activity. To enable the sensitive detection of GVs in intact animals, imaging techniques must overcome the background linear scattering of tissues. This task is accomplished by ultrasound pulse sequences, such as amplitude modulation, which exploit the ability of GVs to produce nonlinear ultrasound scattering (7–9). This ability hinges on the mechanical buckling of GVs – an abrupt transition in mechanical response due to an external load. Specifically, above a threshold acoustic pressure known as the buckling pressure, the protein shell of a GV abruptly undergoes mechanical instability by exhibiting large, reversible deformations, which, in turn, lead to nonlinear scattering of ultrasound waves (7, 10, 11). Previous work has shown that the protein composition of the GV shell can affect GV mechanical properties and acoustic buckling behavior (2, 6, 10). However, the effect of GV geometry on buckling mechanics and ultrasound responsiveness remains uncharacterized. Distinct classes of GVs exhibit different characteristic dimensions with respect to length and diameter (12), and the distribution of these parameters can depend on the cell type expressing the GVs (4).

In this work, combining computational modeling and experimental studies, we systematically investigate how the geometry of cylindrical GVs can affect their buckling behavior upon application of ultrasound pressure. Based on the dimensions of wild-type GVs obtained from cryogenic-electron microscopy (cryo-EM), we developed a series of finite element models of GVs, each with a distinct length or diameter. Our computational simulations predict that the diameter, rather than the length, can significantly alter the threshold buckling pressure of GVs under ultrasound. We then aimed to corroborate these predictions through experiments. To this end, we sorted GVs expressed by cyanobacteria into different populations based on diameter and recorded their respective nonlinear ultrasound scattering. We show that GVs with larger diameters exhibit stronger scattering of nonlinear ultrasound signals for a given acoustic pressure. This work reveals a fundamental relationship between GV geometry and buckling behavior, which provides guidance for the engineering of GVs with different sonomechanical characteristics for enhanced ultrasound imaging and potential multiplexed detection (2).

## MATERIAL AND METHODS

### Computational modeling of GV buckling

We developed a finite element model of a single stripped GV (13) isolated from *Anabaena flos-aquae* (AnaS), in which we adopt the GV shape and geometry from a cryo-EM image (Fig. 1). The adopted geometry consists of a cylindrical shell with conical ends. In view of experimental observations (Fig. S1a), we assume a uniform GV diameter within the cylindrical segment of the protein shell. We model the protein wall as a continuum shell with a thickness of 2.4 nm and a shell density of 1350 kg/m^3^ (7, 14). In order to account for the rib-like structure of the gas vesicle wall, we incorporate an elastic anisotropic material model, with elastic moduli across and along the principal axis of the GV of 0.98 GPa and 3.92 GPa, respectively (7). We also assign a Poisson’s ratio of 0.499, which produces the desired incompressibility. While the material parameters are not obtained from direct experimental measurements, the values chosen lie within a range of parameters consistent with those of protein-based biological materials (15). The model is then discretized using shell elements. We subject the exterior and the interior surfaces of the GV to an initial pressure of 101 kPa, modeling both the inner gas pressure and the pressure of the surrounding environment. To prevent rigid body modes, in which the entire GV structure would undergo translations and rotations without any elastic deformation, we subject the vertices at both the top and bottom conical ends of the GV to the zero displacement Dirichlet boundary condition. We first conduct linear buckling analysis (LBA) and solve the corresponding eigenvalue problem to obtain the threshold buckling pressures. Upon the onset of buckling, the soft protein shell undergoes large deformations, which cannot be resolved using linear analysis. We therefore solve for post-buckling configurations using explicit dynamic analysis, which is a particularly powerful technique when a computational problem includes measures of discontinuity, such as buckling, in the solution. For the explicit dynamic analysis, two reasonable assumptions are made to account for the inner gas pressure dynamics. First, we assume an isothermal buckling mechanism. Second, we neglect diffusion of gas across the GV shell, and treat the encapsulated gas as trapped within a GV, since the gas efflux time is substantially longer than an ultrasound cycle at the frequency used in this study (16, 17). Using these assumptions, we simulate the GV response to acoustic excitation by applying an additional oscillatory overpressure in the form of a tapered sine-burst pulse amplitude signal applied for 1 microsecond at a frequency of 11.4 MHz, which is a typical ultrasound setting used experimentally for imaging. Moreover, in the explicit dynamic analysis, we introduce numerical bulk viscosity damping to eliminate numerical artifacts and to smear nonphysical oscillations in the solutions obtained by utilizing linear and quadratic damping coefficients equal to 0.06 and 1.2, respectively. The selected element size in the discretization of the model is at least one-tenth of the dilatational wavelength, and time steps are automatically incorporated to ensure satisfaction of the Courant-Friedrichs-Lewy (CFL) stability criterion (18). Calculations for both linear buckling analysis and explicit dynamic analysis are carried out using Abaqus (Dassault Systèmes Simulia, France).

**Figure 1.**
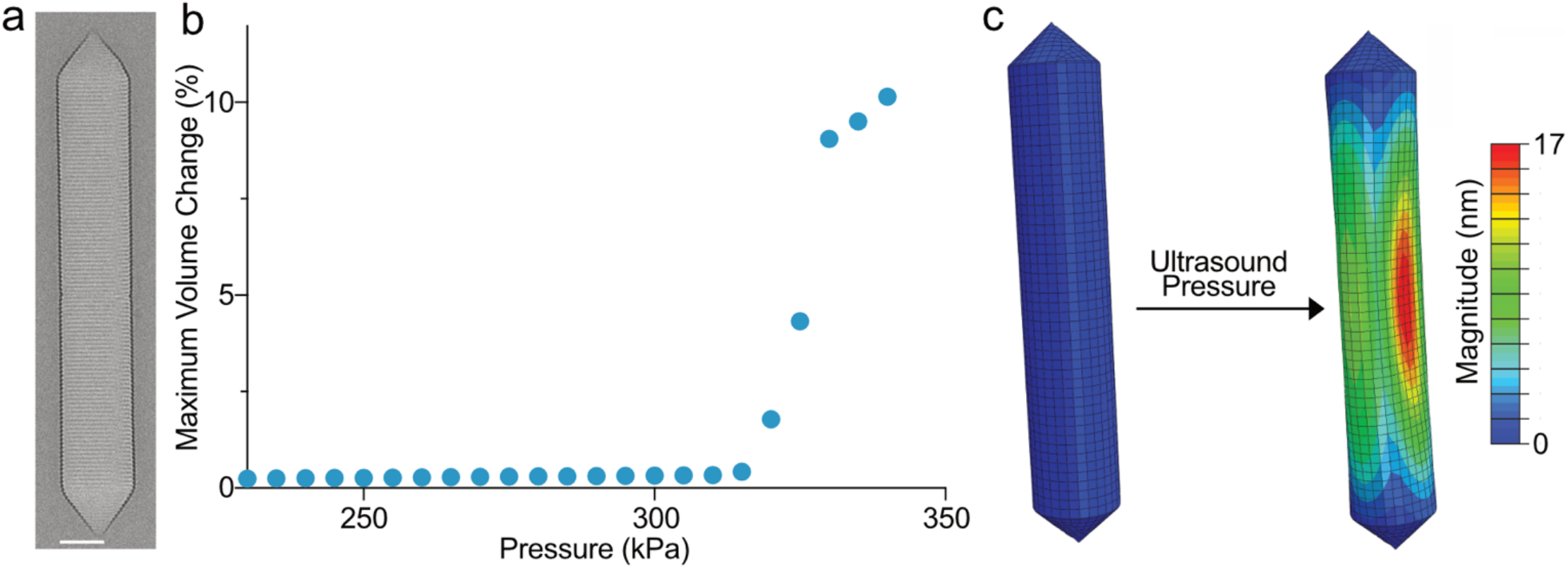
Geometric characterizations and computational modeling of GVs. (a) Representative cryo-EM image of a stripped GV (AnaS) isolated from cyanobacterium *Anabaena flos-aquae*. Scale bar, 50 nm. (b) Maximum percent volume change in a GV as a function of applied ultrasound pressure. The sudden departure from a linear response indicates the onset of buckling in a GV, which is reminiscent of pitch-fork instability. (c) Depictions from a finite element model of a GV with length and diameter dimensions of 500 nm and 85 nm, respectively. Both the initial configuration (left) and the buckled configuration at 331 kPa (right) are depicted.

### GV preparation and quantification

GVs were purified from *Anabaena flos-aquae* as previously described (2, 13). 6 M urea solution was added to purified native GVs and two subsequent rounds of centrifugal flotation and removal of subnatant were preformed to prepare stripped GVs (AnaS). Two rounds of dialysis in PBS were performed to exchange the media. We determined the concentration of GVs by measuring the optical density at 500 nm (OD_500_) with a spectrophotometer (NanoDrop ND-1000, Thermo Scientific). For AnaS, OD_500_ = 1 corresponds to a concentration of 184 pM or a volume fraction of 0.04% of GVs in an aqueous suspension.

### Cryo-EM characterization and image analysis

The geometry of AnaS samples subjected to pre-collapse pressures was characterized using cryo-EM as described before (12). A 3-μL volume of a sample with OD_500_ = ∼5 was applied to C-Flat 2/2-3C grids (Protochips) that were freshly glow-discharged (Pelco EasiGlow, 10 mA, 1 min). GV samples were frozen using a Mark IV Vitrobot (FEI, now Thermo Fisher Scientific) (4°C, 100% humidity, blot force 3, blot time 4 s). Micrographs were collected on a 200 kV Talos Arctica microscope (FEI, now Thermo Fisher Scientific) equipped with a K3 6k × 4k direct electron detector (Gatan). Multi-frame images were collected using SerialEM 3.39 software (19) with a pixel size of 1.17 Å (36,000× magnification) and a defocus of -2.5 μm. Super-resolution movies were corrected for gain reference, binned by a factor of 2, and motion-corrected using MotionCor2 (20). GV dimensions were measured using IMOD 4.12 (21). Statistical analysis was performed in GraphPad PRISM.

### GV diameter consistency analysis

To quantify the stability of the diameter of individual GVs from multiple cryo-EM images, we selected start and end coordinates for individual GVs and subsequently cropped the cylindrical GV tube into segments with ∼10 nm distance using RELION (22). To obtain accurate estimates of GV diameter, we analyzed density profiles for each segment located in the central section of the GV tube using Fiji (23) (Fig. S1a). To evaluate diameter consistency, we calculated the standard deviation of each GV diameter as a percentage of the mean diameter (Fig. S1b).

### Collapse of GVs with defined pressure

A sample of purified AnaS with OD_500_ = ∼20 was loaded in a sealed flow-through quartz cuvette (Hellma Analytics, Plainview, NY) connected to a pressure controller (Alicat Scientific, Tucson, AZ) with N_2_ gas supplied to apply a headspace overpressure. The pressure was slowly increased by 20 kPa at each step, and the OD_500_ was measured with a spectrophotometer (EcoVis, OceanOptics, Winter Park, FL).

### Ultrasound imaging of GVs and image analysis

10 μL of GVs were dispersed in 10 μL of 1% (mass/volume) agarose in PBS and loaded into a homemade gel phantom made of 1% agarose, with a final OD_500_ = 2 measured with a spectrophotometer (NanoDrop ND-1000, Thermo Scientific). A Verasonics Vantage programmable ultrasound scanning system with an L22-14v 128-element linear array transducer (Verasonics, Kirkland, WA) transmitting at 15.6 MHz was used to perform ultrasound imaging. The gel phantom and transducer tip were both immersed in a volume of PBS to conduct imaging. A customized nonlinear ultrasound imaging protocol, namely cross-amplitude modulation (x-AM) (8), was used to specifically characterize the nonlinear contrast of GVs at a distance of 5 mm from the transducer. Specifically, an automated voltage ramp script implemented in MATLAB was used to acquire x-AM signals at each specified voltage step ranging from 1.6 V (corresponding to a peak positive pressure of 150 kPa) to 10 V (corresponding to a peak positive pressure of 734 kPa) with 0.5-V increments. The transmitted pressure level was calibrated using a fiber-optic hydrophone (Precision Acoustics), and the peak positive pressure was termed “acoustic pressure” as shown in Figure 4.

## RESULTS AND DISCUSSION

### Computational analysis

We first investigated the effect of geometric features of GVs on their buckling response under ultrasound. We chose wild-type GVs expressed by cyanobacterium *Anabaena flos-aquae* as a model system (Ana GVs) due to their common use in ultrasound studies. The shell wall of Ana GV is made of GvpA, a primary GV structural protein, and GvpC, a secondary GV structural protein (24). Previous experiments showed that stripped Ana GVs (AnaS), in which GvpC units have been selectively removed or digested, buckle and scatter nonlinearly above a certain acoustic pressure (2, 6–8). As described in the Materials and Methods section for finite element analysis, we modeled the buckling of a stripped GV subjected to ultrasound overpressure. We first conducted simulations using a GV with an average length and diameter of 500 nm and 85 nm, respectively, which correspond to the average dimensions of wild-type Ana GVs (12). We conducted a linear buckling analysis (LBA), in which an eigenvalue problem is formulated upon the construction of the pertinent stiffness and mass matrix. We solved this problem using the Lanczos algorithm and obtained the first ten modes of buckling. Fig. S2 depicts these buckling modes, with the first threshold buckling pressure predicted to occur at 332 kPa. Next, we solved the deformed post-buckling configurations and validated the results of the LBA. The compliant nature of the GV protein shell leads to large deformations upon buckling, which requires nonlinear analysis to resolve. The combination of a compliant protein shell and subsequent nonlinear geometric effects under ultrasound results in an output of ill-conditioned tangent matrices. To compute threshold buckling pressures under these conditions, we utilized a dynamic relaxation approach through explicit analysis. To compute the threshold buckling pressures for each buckling mode obtained, we conducted a series of simulations, independent of the LBA analysis, for an individual stripped GV, where the overpressure varies over a period of 1 microsecond, starting at 100 kPa and increasing in steps of 20 kPa until a pressure that causes the structure to buckle. Each simulation was designed with a total simulation time of 1 microsecond at 11.4 MHz frequency.

We quantified GV deformations by measuring the change in volume, which, prior to the onset of buckling, increases negligibly with externally applied cycles of ultrasound pressure. At the threshold pressure for the onset of buckling, an abrupt transition occurs in the GV deformation mechanics. Notably, this transition may not occur in response to all the cycles within an ultrasound pulse, due to the tapered nature of pulse amplitudes and to the nonlinear geometric effects of a GV exposed to ultrasound, which may induce the onset of buckling only after the GV experiences a few cycles of ultrasound pressure. We then identified the exact threshold buckling pressure that causes this nonlinear response, within a narrow range of 1 kPa, via a bisection method in which we ascertain the interval that contains the threshold buckling pressure by repeatedly bisecting each pressure interval and selecting the subinterval in which buckling commences. This bisection algorithm determined the threshold buckling pressure of AnaS GVs with dimensions of 500 nm in length and 85 nm in diameter to be 331 kPa (Fig. 1).

Notably, it is possible that accounting for nonlinear deformations using explicit dynamic analysis based on volumetric changes may lead to a threshold buckling pressure lower than that obtained via LBA. These nonlinear deformations can accommodate buckling at pressures below the values obtained from LBA. We also conducted several numerical tests covering a range of mesh sizes and verified that the results of our calculations were not affected by the discretization resolution (Fig. S3).

### Geometry-dependent GV buckling

After validating a computational model that captures the ultrasound-induced buckling of a GV with fixed geometry, we aimed to model the effect of different GV lengths and diameters on the threshold buckling pressure. To this end, we conducted a thorough sensitivity analysis using an exhaustive search approach, in which we created several distinct computational GV models, each of which having identical material properties, boundary conditions, and loading conditions, including an identical ultrasound pressure waveform. In these computational models, we also fixed the GV length (or diameter) and varied the GV diameter (or length) across a physiologically relevant range of values (12, 13). In each of the corresponding finite element models, the element type and the mesh size remained invariant, leading to a different number of elements and nodes across models. Additionally, for each model, with the details delineated in the previous subsection, both LBA and explicit dynamic analysis were conducted.

We first investigated the dependence of threshold buckling pressures on GV diameter. We created two sets of models with fixed GV lengths of 300 nm or 500 nm, and in each set of models, we simulated a physiologically relevant range of GV diameters (12, 13). The results of the simulations are shown in Fig. 2 with representative snapshots of the buckled GV configuration, which demonstrates that varying the GV diameter substantially impacts the threshold buckling pressure value. We quantified this dependency using a curve fit that is defined as *P* = *AD*^*α*^ + *B*, with *P* and *D* being the buckling pressure and the GV diameter, respectively, *A* and *B* being fitting parameters. We consequently obtained a value of *α* ≅ −3. Next, we investigated the dependence of threshold buckling pressures on GV length. Fig. 3 shows the results of simulations conducted for two distinct GV diameters: 60 nm and 83 nm, with illustrative depictions of buckled configurations for three representative GVs. In dramatic contrast to our results with varying diameter, the length sensitivity analysis shows that the threshold buckling pressure is virtually unaffected by differences in GV length over the typical range exhibited by AnaS GVs. This result is apparent with the exponent *α* being close to zero, obtained by fitting a function of the form *P* = *AL*^*α*^ + *B* to the data, where *P* and *L* are the threshold buckling pressure and the GV length, respectively.

**Figure 2.**
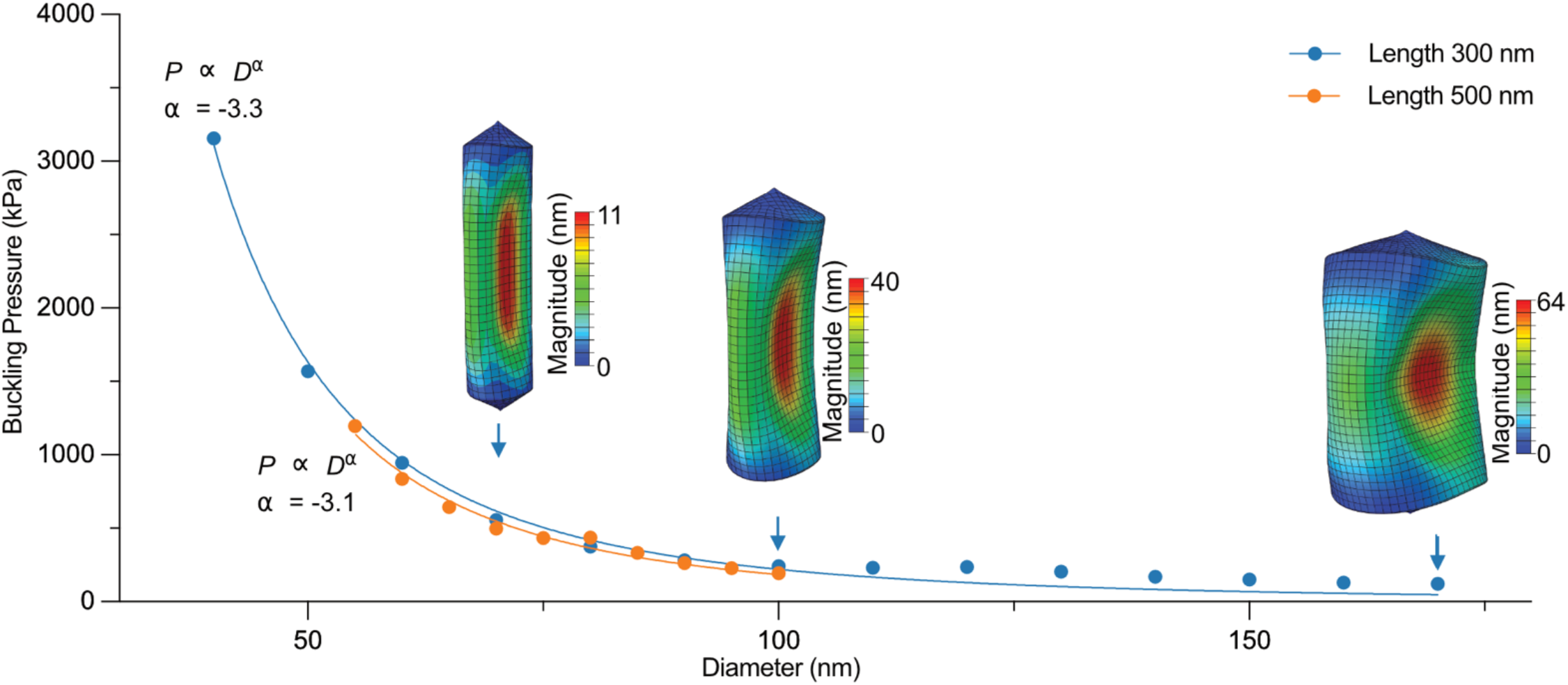
Diameter sensitivity analysis of GV buckling. The effect of GV diameter on the threshold buckling pressure at two fixed GV lengths: 300 nm (blue) and 500 nm (orange). Diagrams from simulations illustrate the buckled configuration of GVs with a fixed length of 300 nm and different diameters of 70 nm (*left*), 100 nm (*middle*), and 170 nm (*right*).

**Figure 3.**
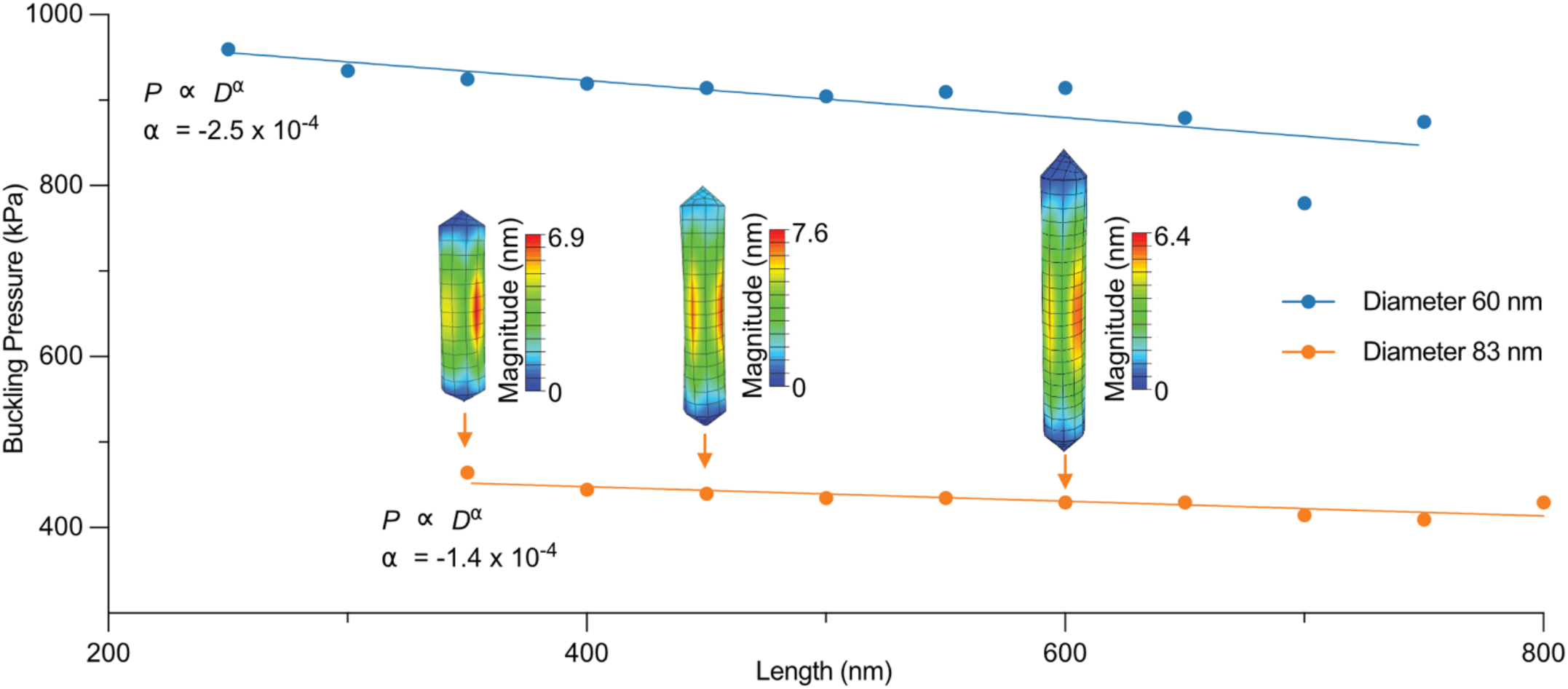
Length sensitivity analysis of GV buckling. The effect of GV length on the threshold buckling pressure at two fixed GV diameters: 60 nm (blue) and 83 nm (orange). Diagrams from simulations illustrate the buckled configurations of GVs with a fixed diameter of 83 nm and different lengths of 350 nm (*left*), 450 nm (*middle*), and 600 nm (*right*).

Considering the unsubstantial effect of GV length on the onset of buckling, we remark that the theory of cylindrical shells can help interpret our results for GV buckling. By examining the buckling theory of a shell subjected to external pressure, as well as the solutions of the corresponding eighth-order governing differential equation (also known as the Donnell stability equation), we determined the results for the modulus, Poisson’s ratio, and the moment of inertia of the cross section (25). Although our computational models of GVs account for an anisotropic finite-length shell with conical ends, the agreement of the integer parts of the exponents obtained from our simulations with those obtained from the idealized shell theory further posits that the diameter is the dominant dimensional feature influencing GV buckling.

### Experimental validation

To experimentally validate the geometry-buckling relationship revealed by our simulation results, we first fractionated AnaS into different size distributions. Given that these GVs are expressed in a single species of cyanobacteria harboring the same gene cluster, we assume the material properties of the major structural protein of the shell (GvpA) to be the same and not dependent on GV geometry. To obtain a different size distribution of AnaS, we slowly increased the hydrostatic pressure around AnaS, leading to the irreversible collapse of some GVs, and characterized the geometry of the remaining GVs. Since previous studies showed a correlation between the threshold limiting case of *L* >> *D*. For an isotropic shell, it can be shown that the buckling pressure *P* satisfies 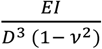, with *E, v*, and *I* representing the Young acoustic buckling pressure of a GV and its hydrostatic collapse pressure (2, 6), our simulation results led us to hypothesize that GVs surviving higher pressures without collapse would have smaller diameters and generate less buckling-induced nonlinear ultrasound collapse pressure than the original GV population, indicating that our hydrostatic pressure treatment at 200 contrast. Because collapsed GVs do not scatter light as do intact GVs, we quantified the number of intact GVs remaining by measuring the optical density at a wavelength of 500 nm (OD_500_) after exposure to different hydrostatic pressures. The OD_500_ of AnaS remained unchanged when exposed to low hydrostatic pressure, and it significantly decreased above a certain pressure until all GVs collapsed. By setting the applied hydrostatic pressure at 200 kPa, approximately 30% of GVs remain intact, whereas at a pressure of 220 kPa, only ∼10% GVs remain intact (Fig. 4A). Notably, GVs that remain intact at 200 kPa have a higher hydrostatic kPa successfully selected GVs that are mechanically more robust and resist higher hydrostatic pressures. absolute length and diameter distributions of pressure-treated GV samples were characterized by cryo-EM (Fig. 4B). We found that the length distribution of GVs does not change significantly, and is independent of pressure treatment (Fig. 4C). However, it is clear that increasing applied hydrostatic pressure led to smaller average diameters in remaining intact GVs (Fig. 4D). Specifically, a 200 kPa pre-collapse step destroys any GVs with a diameter larger than 90 nm. This observation supports the prediction that the mechanical properties of GVs depend on the diameter – but not length, resulting in significantly different susceptibility to hydrostatic pressure.

**Figure 4.**
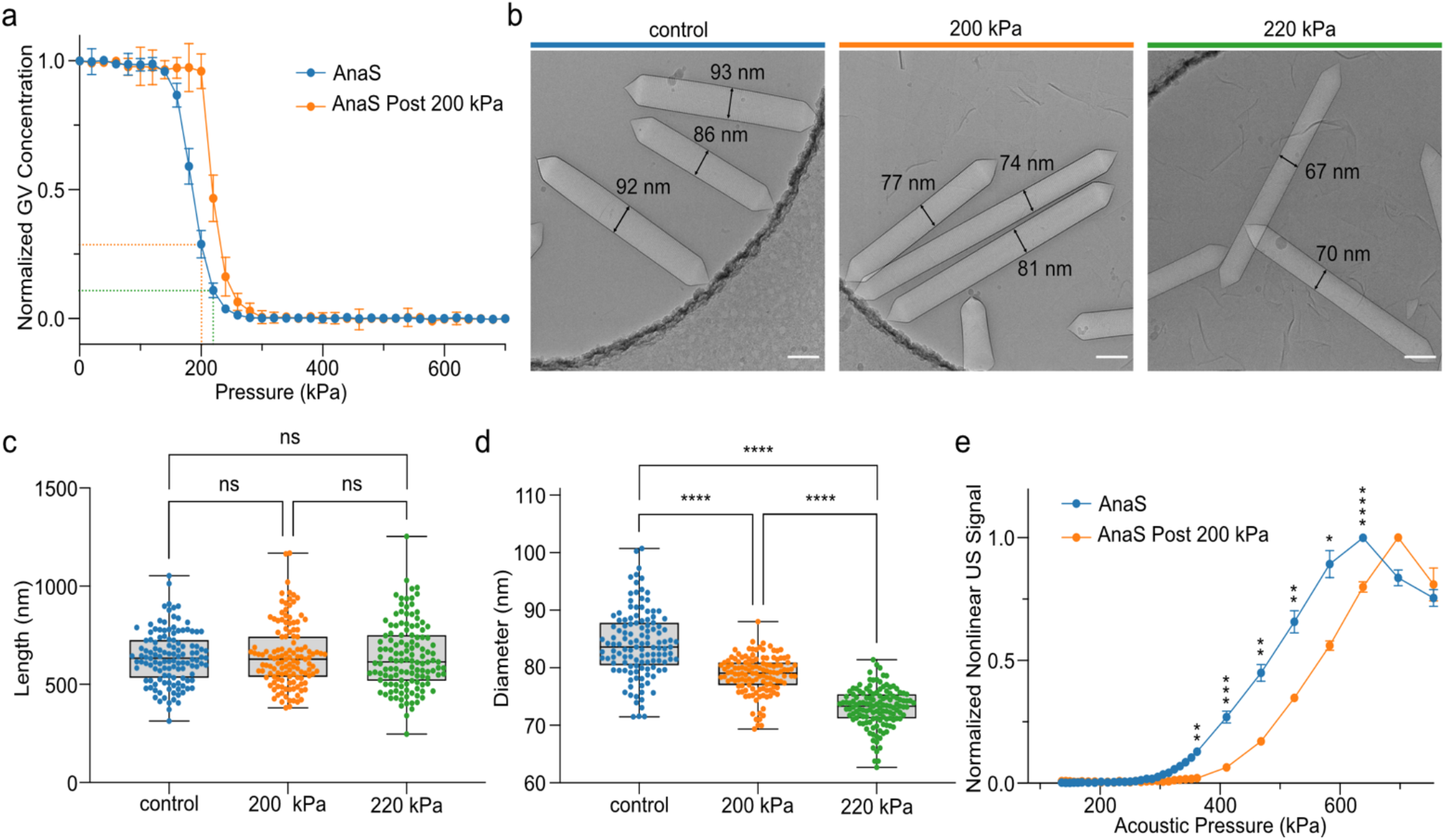
Experimental validation of the GV geometry-buckling relationship. (a) Hydrostatic collapse pressure curves for stripped GVs isolated from cyanobacterium Anabaena flos-aquae (AnaS), with (orange) and without (blue) pre-collapse hydrostatic pressure treatment at 200 kPa. Dashed lines indicate the pressure and corresponding OD500 for samples that were collected for cryo-EM and nonlinear ultrasound imaging analysis. (b) Representative cryo-EM images of AnaS used to measure lengths and diameters of GVs from the control sample (left), and after incubation at a hydrostatic pressure of 200 kPa (middle) and 220 kPa (right). Scale bars, 100 nm. (c) Length and (d) diameter distributions of the intact GV fraction after exposure to the indicated hydrostatic pressure. Asterisks indicate statistical significance by one-way ANOVA tests (**** = p < 0.0001); ns = no significance. (e) Nonlinear ultrasound signals from AnaS (n = 4) as a function of acoustic pressure from samples with (orange) and without (blue) pre-collapse hydrostatic pressure treatment at 200 kPa. Asterisks represent statistical significance by unpaired t-tests (**** = p < 0.0001, *** = p <0.001, ** = p <0.01, * = p <0.05).

The sonomechanical buckling behavior of GVs with different size distributions was studied using nonlinear ultrasound imaging, which detects nonlinear scattering signals generated by GV buckling. As hydrostatic pressure treatment of GV samples at 220 kPa reduces the number of GVs below the level needed for reliable ultrasound imaging, we proceeded with imaging only AnaS without pre-treatment or after pre-treatment with 200 kPa. We found that, at the same concentration of intact GVs, pressure-treated GVs require a higher threshold pressure to generate detectable nonlinear signal compared to GVs that did not undergo pre-collapse treatment (Fig. 4E). This set of results agrees well with our modeling prediction that GVs with larger diameters buckle at lower threshold pressures and would thus be expected to generate nonlinear signals at lower pressures compared to GVs with smaller diameters. The apparent experimental buckling thresholds were 300 kPa and 350 kPa for untreated AnaS and pre-collapsed AnaS, respectively. These experimental values are not far from the threshold buckling pressures predicted by our model (263 kPa and 331 kPa, respectively) based on the largest diameter observed in a sample population of GVs, supporting the general validity of our simulations. The fact that our experimental values for threshold buckling pressure are slightly larger than the computationally predicted values can be explained by the fact that only a small fraction of GVs possess the largest diameter observed in a given sample population, and the sample may therefore not generate a detectable amount of ultrasound signal until GVs with smaller diameters start to buckle at higher pressures. Notably, the pressure pre-treated GV sample exhibited a peak nonlinear ultrasound signal at a higher pressure (above which the signal declines due to acoustic collapse of the GVs) than the GV sample not subjected to pre-collapse treatment, again suggesting that the pressure required to collapse GVs becomes higher after pre-collapse treatment. Experimental validation further supports the correlation between the hydrostatic collapse pressure and threshold acoustic buckling pressure: GVs with lower hydrostatic collapse pressures tend to buckle at lower acoustic pressures and generate higher cross-amplitude modulation (x-AM) signal than GVs with higher hydrostatic collapse pressures under the same ultrasound conditions, a result which has also been observed in other studies (2, 6).

## CONCLUSION

The sonomechanical buckling properties of GVs were systematically investigated through finite element simulations and experiments. Computational results predicted that the GV diameter, but not the length, strongly influences the buckling behaviors of GV. We have determined that there is an inverse cubic relation between the threshold buckling pressure and the GV diameter. Above the threshold buckling pressure, ultrasound is predicted to induce large deformations of the GV shell, which agrees with the experimentally observed nonlinear acoustic backscattering response of GVs. Our computational models and analysis were corroborated by the results of experiments using nonlinear ultrasound imaging of GVs having the same genotype but different size distributions. Our results elucidate the effect of geometry on the sonomechanical buckling of GVs, which has the potential to guide future engineering of GVs as highly sensitive and specific ultrasound contrast agents, reporter genes and biosensors, the advancement of high-precision, nonlinear imaging. In addition, mechanical insights into GV interactions with ultrasound waves may benefit other GV-enabled technologies such as acoustic manipulation of engineered cells and cell-based therapeutics (26, 27).

## ACKNOWLEDGMENTS

The authors are grateful to Ngozi A. Eze for the helpful editorial comments. This research was supported by the National Institutes of Health grant R01-EB018975. Related research in the Shapiro Lab is supported by the Packard Foundation, The Pew Charitable Trusts, and the Chan Zuckerberg Initiative. Cryo-electron microscopy was performed at the Beckman Institute Resource Center for Transmission Electron Microscopy at Caltech. M.G.S. is an Investigator of the Howard Hughes Medical Institute (HHMI).

This article is subject to HHMI’s Open Access to Publications policy. HHMI Investigators have previously granted a nonexclusive CC BY 4.0 license to the public and a sublicensable license to HHMI in their research articles. Pursuant to those licenses, the author-accepted manuscript of this article can be made freely available under a CC BY 4.0 license immediately upon publication.

## AUTHOR CONTRIBUTIONS

H.S., Y.Y., P.D., and M.G.S. conceived and designed the study. H.S. and E.M. developed the computational models. H.S. and E.M. performed the simulations and analyzed the simulation data. Y.Y., P. D., Z. J., and D. M. conducted *in vitro* experiments and analyzed the experimental data. N.N.N. was involved in planning experiments and data analysis. M.G.S., M.O., and G.J.J. supervised the research. H.S., Y.Y., P.D., and M.G.S. wrote and edited the manuscript. All authors read, edited, and confirmed the content of the manuscript.

## SUPPLEMENTARY MATERIALS

**Figure S1.**
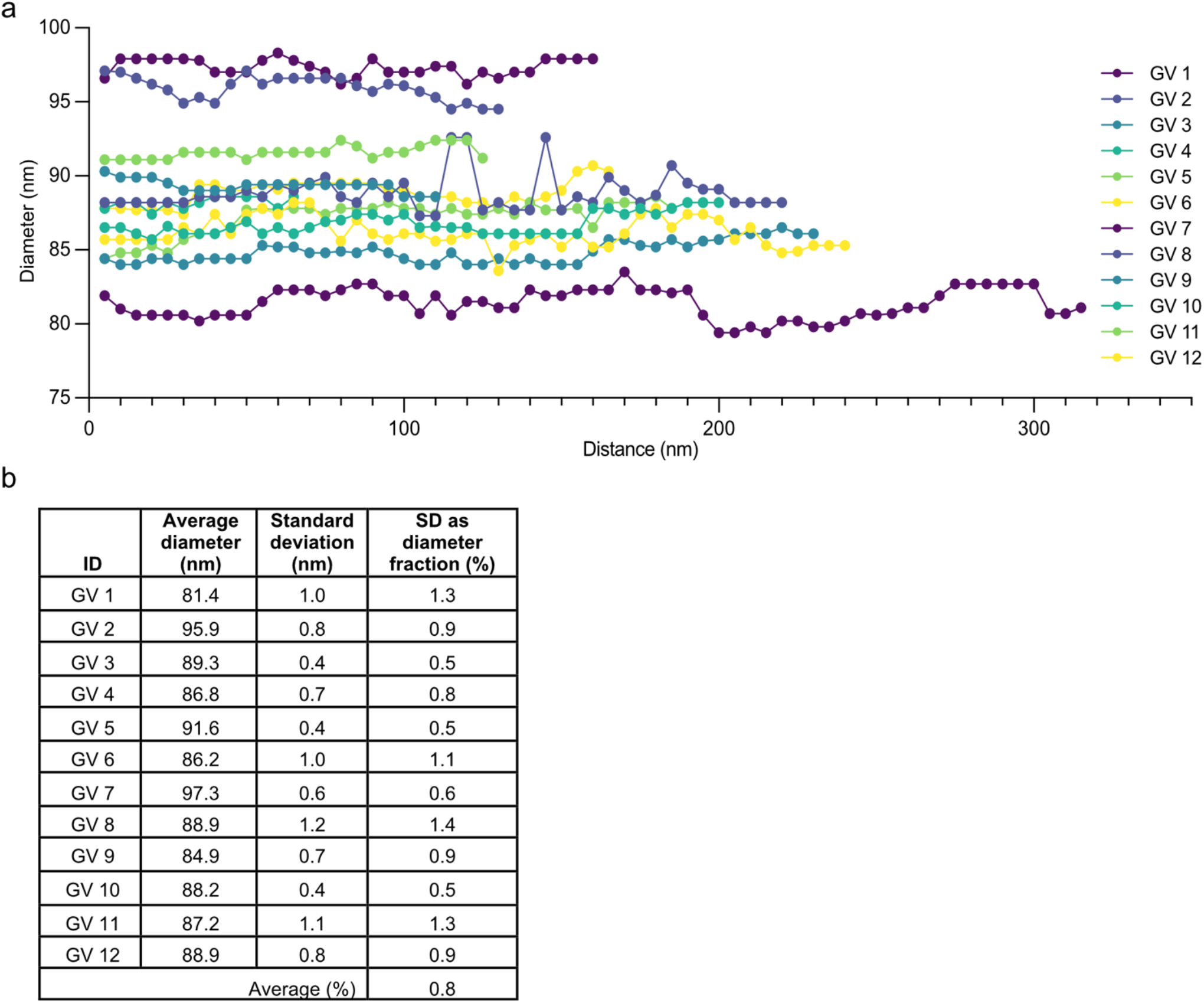
Diameter consistency analysis. (a) Diameter measurements of individual GVs. For each individual GV, the diameter was measured at increments of 10 nm along the main axis. (b) Table of summarized data showing diameter consistency across individual GVs.

**Figure S2.**
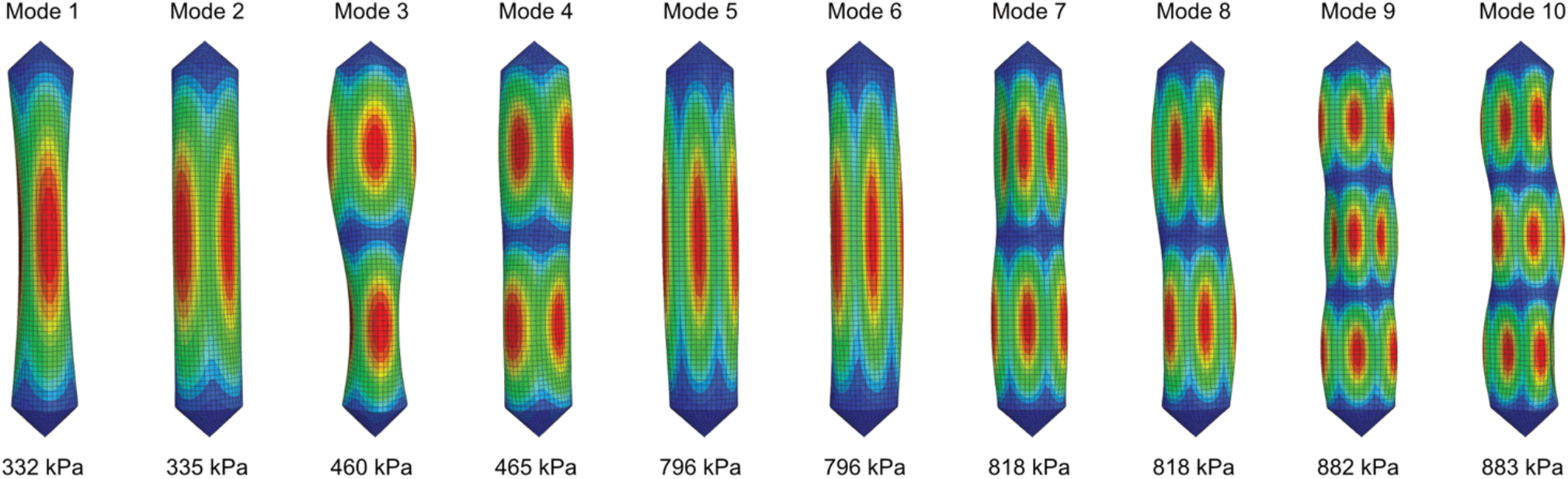
The first ten modes of buckling (*i*.*e*., eigenvectors) and the corresponding threshold buckling pressures (*i*.*e*., eigenvalues) obtained through linear buckling analysis (LBA).

**Figure S3.**
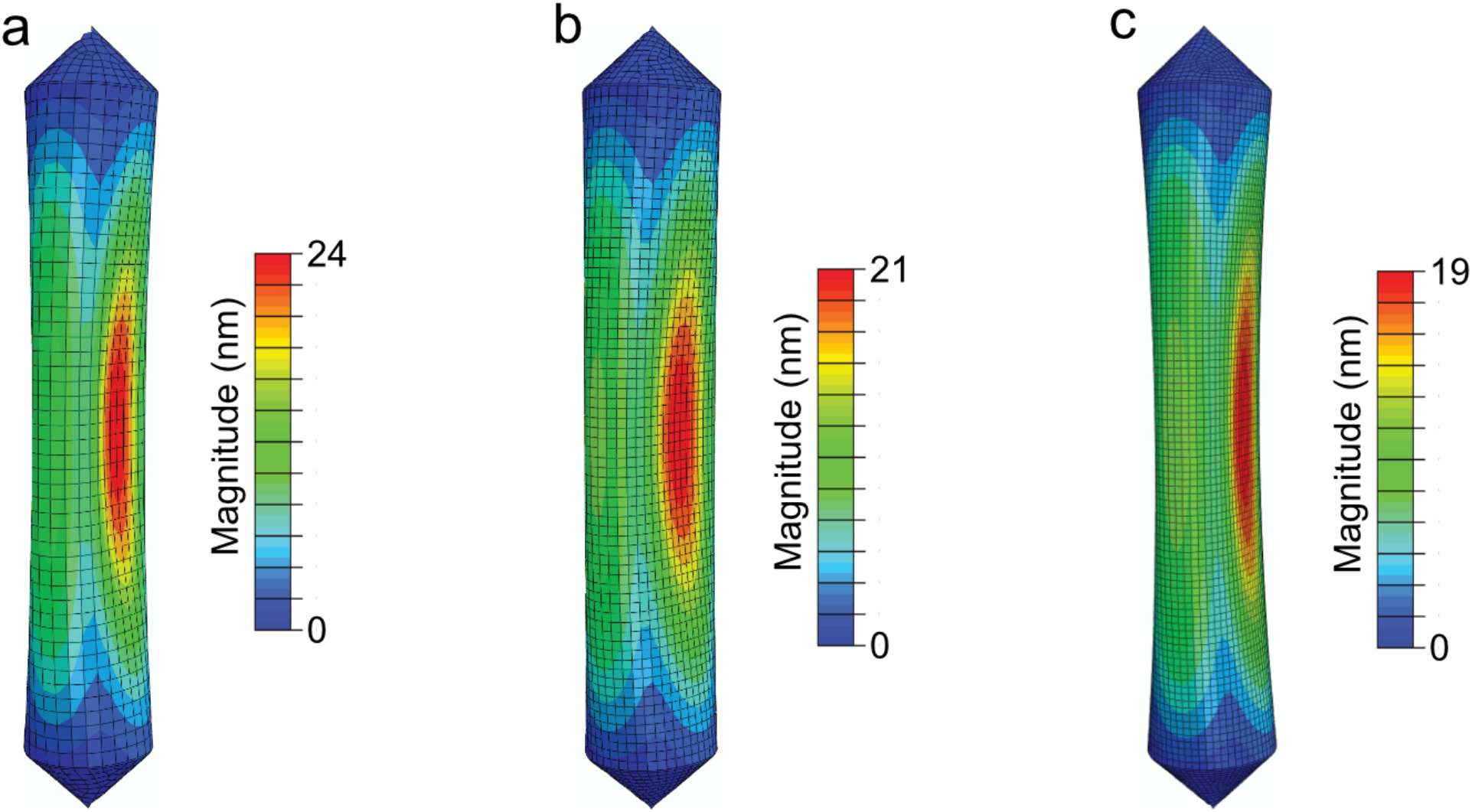
Mesh sensitivity analysis. GV buckling simulations (with GV length and diameter set to 500 nm and 85 nm, respectively) are carried out using three different mesh sizes. A buckled configuration is depicted for each of the three models with discretization lengths as follows: (a) 9 nm, (b) 6 nm, and (c) 4.5 nm. Our results show that all three models predict an identical threshold buckling pressure of 331 kPa.

## REFERENCES

1. Shapiro, M.G., P.W. Goodwill, A. Neogy, M. Yin, F.S. Foster, D.V. Schaffer, and S.M. Conolly. 2014. Biogenic gas nanostructures as ultrasonic molecular reporters. Nat. Nanotechnol. 9:311–316.

2. Lakshmanan, A., A. Farhadi, S.P. Nety, A. Lee-Gosselin, R.W. Bourdeau, D. Maresca, and M.G. Shapiro. 2016. Molecular Engineering of Acoustic Protein Nanostructures. ACS Nano. 10:7314–7322.

3. Bourdeau, R.W., A. Lee-Gosselin, A. Lakshmanan, A. Farhadi, S.R. Kumar, S.P. Nety, and M.G. Shapiro. 2018. Acoustic reporter genes for noninvasive imaging of microorganisms in mammalian hosts. Nature. 553:86–90.

4. Farhadi, A., G.H. Ho, D.P. Sawyer, R.W. Bourdeau, and M.G. Shapiro. 2019. Ultrasound imaging of gene expression in mammalian cells. Science. 365:1469–1475.

5. Hurt, R.C., M.T. Buss, M. Duan, K. Wong, M.Y. You, D.P. Sawyer, M.B. Swift, P. Dutka, D.R. Mittelstein, Z. Jin, M.H. Abedi, A. Farhadi, R. Deshpande, and M.G. Shapiro. Genomically Mined Acoustic Reporter Genes Enable In Vivo Monitoring of Tumors and Tumor-Homing Bacteria.

6. Lakshmanan, A., Z. Jin, S.P. Nety, D.P. Sawyer, A. Lee-Gosselin, D. Malounda, M.B. Swift, D. Maresca, and M.G. Shapiro. 2020. Acoustic biosensors for ultrasound imaging of enzyme activity. Nat. Chem. Biol.

7. Maresca, D., A. Lakshmanan, A. Lee-Gosselin, J.M. Melis, Y.-L. Ni, R.W. Bourdeau, D.M. Kochmann, and M.G. Shapiro. 2017. Nonlinear ultrasound imaging of nanoscale acoustic biomolecules. Appl. Phys. Lett. 110:073704.

8. Maresca, D., D.P. Sawyer, G. Renaud, A. Lee-Gosselin, and M.G. Shapiro. 2018. Nonlinear X-wave ultrasound imaging of acoustic biomolecules. Phys Rev X. 8.

9. Rabut, C., D. Wu, B. Ling, Z. Jin, D. Malounda, and M.G. Shapiro. 2021. Ultrafast amplitude modulation for molecular and hemodynamic ultrasound imaging. Appl. Phys. Lett. 118:244102.

10. Cherin, E., J.M. Melis, R.W. Bourdeau, M. Yin, D.M. Kochmann, F.S. Foster, and M.G. Shapiro. 2017. Acoustic Behavior of Halobacterium salinarum Gas Vesicles in the High-Frequency Range: Experiments and Modeling. Ultrasound Med. Biol. 43:1016–1030.

11. Zhang, S., A. Huang, A. Bar-Zion, J. Wang, O.V. Mena, M.G. Shapiro, and J. Friend. 2020. The vibration behavior of sub-micrometer gas vesicles in response to acoustic excitation determined via laser Doppler vibrometry. Adv. Funct. Mater. 30:2000239.

12. Dutka, P., D. Malounda, L.A. Metskas, S. Chen, R.C. Hurt, G.J. Lu, G.J. Jensen, and M.G. Shapiro. 2021. Measuring gas vesicle dimensions by electron microscopy. Protein Sci. 30:1081–1086.

13. Lakshmanan, A., G.J. Lu, A. Farhadi, S.P. Nety, M. Kunth, A. Lee-Gosselin, D. Maresca, R.W. Bourdeau, M. Yin, J. Yan, C. Witte, D. Malounda, F.S. Foster, L. Schröder, and M.G. Shapiro. 2017. Preparation of biogenic gas vesicle nanostructures for use as contrast agents for ultrasound and MRI. Nat. Protoc. 12:2050– 2080.

14. Pfeifer, F. 2012. Distribution, formation and regulation of gas vesicles. Nat. Rev. Microbiol. 10:705–715.

15. Gosline, J., M. Lillie, E. Carrington, P. Guerette, C. Ortlepp, and K. Savage. 2002. Elastic proteins: biological roles and mechanical properties. Philos. Trans. R. Soc. Lond. B Biol. Sci. 357:121–132.

16. Walsby, A.E., N.P. Revsbech, and D.H. Griffel. 1992. The gas permeability coefficient of the cyanobacterial gas vesicle wall. J. Gen. Microbiol. 138:837–845.

17. Kunth, M., G.J. Lu, C. Witte, M.G. Shapiro, and L. Schröder. 2018. Protein Nanostructures Produce Self-Adjusting Hyperpolarized Magnetic Resonance Imaging Contrast through Physical Gas Partitioning. ACS Nano. 12:10939–10948.

18. Achenbach, J. 2012. Wave Propagation in Elastic Solids. Elsevier.

19. Mastronarde, D.N. 2005. Automated electron microscope tomography using robust prediction of specimen movements. J. Struct. Biol. 152:36–51.

20. Zheng, S.Q., E. Palovcak, J.-P. Armache, K.A. Verba, Y. Cheng, and D.A. Agard. 2017. MotionCor2: anisotropic correction of beam-induced motion for improved cryo-electron microscopy. Nat. Methods. 14:331–332.

21. Kremer, J.R., D.N. Mastronarde, and J.R. McIntosh. 1996. Computer visualization of three-dimensional image data using IMOD. J. Struct. Biol. 116:71–76.

22. Zivanov, J., T. Nakane, B.O. Forsberg, D. Kimanius, W.J. Hagen, E. Lindahl, and S.H. Scheres. 2018. New tools for automated high-resolution cryo-EM structure determination in RELION-3. Elife. 7.

23. Schindelin, J., I. Arganda-Carreras, E. Frise, V. Kaynig, M. Longair, T. Pietzsch, S. Preibisch, C. Rueden, S. Saalfeld, B. Schmid, J.-Y. Tinevez, D.J. White, V. Hartenstein, K. Eliceiri, P. Tomancak, and A. Cardona. 2012. Fiji: an open-source platform for biological-image analysis. Nat. Methods. 9:676–682.

24. Walsby, A.E. 1994. Gas vesicles. Microbiol. Rev. 58:94–144.

25. Timoshenko, S., and S. Woinowsky-Krieger. 1959. Theory of plates and shells. McGraw-hill New York.

26. Wu, D., D. Baresch, C. Cook, D. Malounda, D. Maresca, M.P. Abundo, D.R. Mittelstein, and M.G. Shapiro. 2019. Genetically encoded nanostructures enable acoustic manipulation of engineered cells. 691105.

27. Bar-Zion, A., A. Nourmahnad, D.R. Mittelstein, S. Shivaei, S. Yoo, M.T. Buss, R.C. Hurt, D. Malounda, M.H. Abedi, A. Lee-Gosselin, M.B. Swift, D. Maresca, and M.G. Shapiro. 2021. Acoustically triggered mechanotherapy using genetically encoded gas vesicles. Nat. Nanotechnol. 16:1403–1412.

